# OpenIrisDPI: An Open-Source Digital Dual Purkinje Image Eye Tracker for Visual Neuroscience

**DOI:** 10.1101/2025.04.18.649589

**Authors:** Ryan A. Ressmeyer, Jorge Otero-Millan, Gregory D. Horwitz, Jacob L. Yates

## Abstract

**Background:** Video-based eye trackers are widely used in vision science, psychology, clinical assessment, and neurophysiology. Many such systems track the pupil center and corneal reflection (P-CR) and compare their positions to estimate the direction of gaze. However, P-CR eye trackers are often too imprecise for applications with stringent eye tracking quality requirements.

**New method:** We present OpenIrisDPI, an open-source plugin for the OpenIris framework that implements dual Purkinje image (DPI) tracking. OpenIrisDPI supports simultaneous pupillography, a technique widely used in perceptual psychology and neuroscience, and it enables direct comparison between P-CR and DPI signals.

**Results:** Data collected from macaque monkeys using OpenIrisDPI show that the P-CR method overestimates the amount of fixational drift between saccades compared to DPI. The accuracy of the DPI signal was further validated using high-density extracellular recording of neurons in the lateral geniculate nucleus. Compensating for the effects of fixational eye movements using DPI signals produced sharper estimates of neuronal receptive fields than using simultaneously collected P-CR signals.

**Comparison with existing methods:** OpenIrisDPI is provided as open-source software and operates on consumer-grade hardware, making it more accessible than previously described DPI eye trackers and less costly than many P-CR systems. To our knowledge, OpenIrisDPI is the first eye tracker to perform both pupillography and DPI eye tracking.

**Conclusion:** OpenIrisDPI makes high-precision eye tracking readily available to the research community. It is well suited for visual neuroscience applications, where accurate knowledge of the retinal image during experiments is critical.

**Graphical Abstract:** 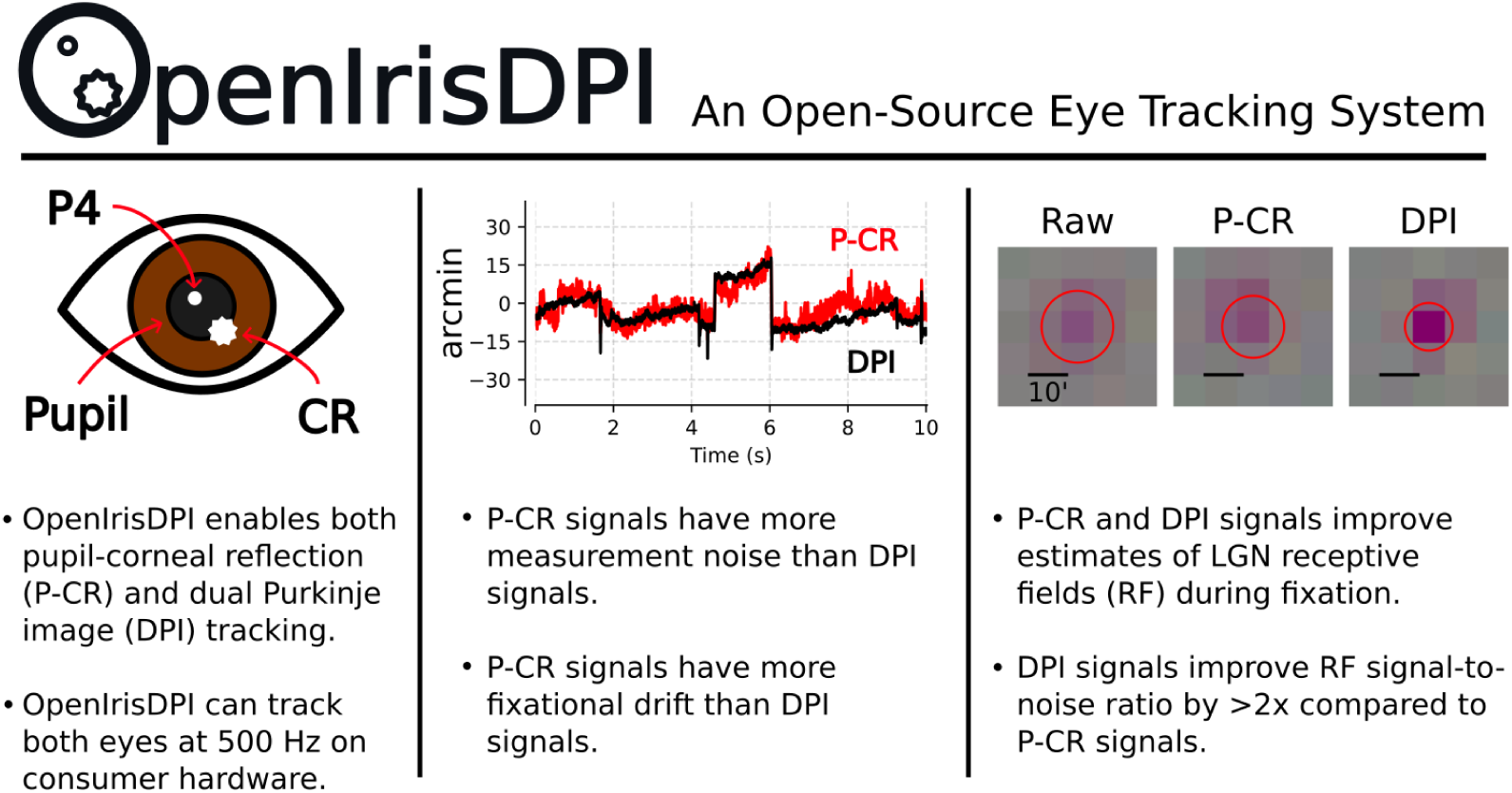

**Highlights:**

- OpenIrisDPI is a new open-source eye tracking system.
- OpenIrisDPI tracks the pupil, corneal reflection, & fourth Purkinje image at 500 Hz.
- Dual Purkinje image-based eye tracking is more precise than pupilbased tracking.
- DPI improves receptive field characterization of LGN neurons in fixating macaques.

## 1. Introduction

Video-based eye tracking has become commonplace in vision research, yet many high-precision applications, from mapping receptive fields to studying fixational eye movements, require measuring the direction of gaze with a precision of a few arcminutes or less (Ko et al. 2016; Yates et al. 2023; see Appendix A for a discussion on precision in the context of eye tracking). Despite the proliferation of video-based eye trackers, many modern systems fail to meet this stringent requirement (Holmqvist and Blignaut 2020). Their performance is often constrained by algorithmic noise and biological artifacts that can introduce errors an order of magnitude larger than the very eye movements—such as microsaccades and ocular drift—they are intended to measure (I. T. Hooge, Hessels, et al. 2019; Wang et al. 2017).

A common method that exemplifies these limitations is pupil-center/corneal-reflection (P-CR) tracking. In this system, a digital camera is used to image the eye, and a computer is used to estimate the positions of the pupil center and corneal reflection. The output of the tracker is the vector difference between these two features—a quantity that is minimally affected by translation of the head relative to the camera (which is difficult to avoid unless the camera is fixed to the head; Guestrin and Eizenman 2006). This invariance to translations of the head is a major advantage of P-CR systems compared to systems that track single features (e.g. pupil-only trackers).

Nevertheless, P-CR systems suffer from two distinct sources of imprecision that limit their utility for high-precision applications. The first is temporally uncorrelated frame-to-frame error in the estimated positions of the pupil center and the CR (measurement noise). The amplitude of measurement noise is often high enough to obscure small eye movements (Niehorster et al. 2021), but is reducible by optimizing algorithms, optics, and imaging hardware (Ivanchenko et al. 2021). The second, more fundamental source of imprecision is pupil decentration, which results from asymmetric movements of the iris during dilation and contraction. These movements cause the pupil center to translate by as much as 0.1 mm between extreme dilation states (Wyatt 1995), which P-CR systems misinterpret as eye movements of a degree or more (Wildenmann and Schaeffel 2013; Wyatt 2010). The pupilary decentration artifact is present in widely used systems, like the EyeLink 1000 and Tobii Pro Spectrum (I. T. Hooge, Niehorster, et al. 2021), and is an inherent consideration for any method that relies on the pupil center.

An alternative optical technique is the dual Purkinje image (DPI) method (Fig. 1). DPI tracking is conceptually similar to P-CR, but it replaces the pupil center with a reflection from the posterior surface of the lens (the fourth Purkinje image, or P4). This method shares the head translation-invariance of P-CR and has the additional advantage of insensitivity to pupil decentration.

**Figure 1:**
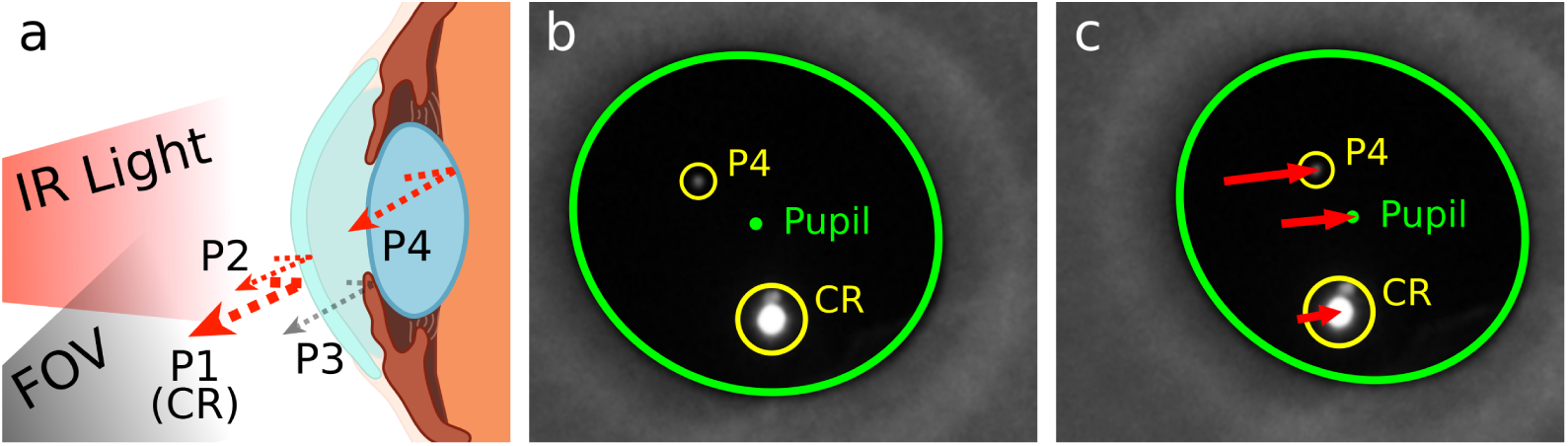
Differential motion of features during eye rotation enables DPI and P-CR tracking. a) Anatomical cross-section showing the four optical surfaces that generate Purkinje reflections: outer cornea (P1 or CR), inner cornea (P2), outer lens (P3), and inner lens (P4). The eye is illuminated with an infrared (IR) light, and reflections are within a camera’s field of view (FOV). DPI and P-CR eye trackers are distinguished by the use of either P4 or the pupil center. Both methods track the CR. b) Images of the eye showing the CR and P4 (marked in yellow), with the pupil contour fitted by a green ellipse. The center of the pupil ellipse is depicted by a green dot and labeled. c) Image captured after a leftward saccade, with red arrows indicating the displacement of CR, P4, and pupil center from their positions in (b). Note that the CR, P4 and the pupil center move by different amounts during eye rotation but would move together during head translation, allowing eye trackers to distinguish between these movements.

Historically, DPI systems relied on custom optoelectronic feedback loops that were expensive, technically complex, and difficult to operate (Crane and Steele 1985). This limited their use to a handful of specialized labs. With the recent development of video-based methods for tracking the Purkinje reflections (Abdulin et al. 2019; Lu et al. 2020; Tabernero and Artal 2014; Wu et al. 2022), digital DPI tracking is poised to overcome these barriers to broader adoption.

We present OpenIrisDPI with the goal of making high-precision DPI eye tracking widely available to the research community (Fig. 2). OpenIrisDPI is an eye tracking system built using the OpenIris eye tracking framework (Sadeghi et al. 2024) that delivers several novel features: (1) simultaneous pupil and DPI tracking, (2) 500 Hz binocular sampling using modern consumer CPUs, and (3) an optional oblique imaging configuration that eliminates the need for hot mirrors. Unlike most conventional P-CR based eye trackers, OpenIrisDPI can measure pupilary dynamics and precise gaze position simultaneously and without conflation. The source code for the plugin is free and openly available under the GPLv3 license (https://github.com/ryan-ressmeyer/OpenIrisDPI). The associated wiki provides an overview of DPI principles and analysis techniques, as well as instructions for building imaging systems, setting parameters, integrating OpenIris with experimental systems, and synchronizing recorded timestamps with independent data streams. At the time of publication, a custom digital DPI system with sub-arcminute measurement noise (see Results) can be assembled for less than $5000.

**Figure 2:**
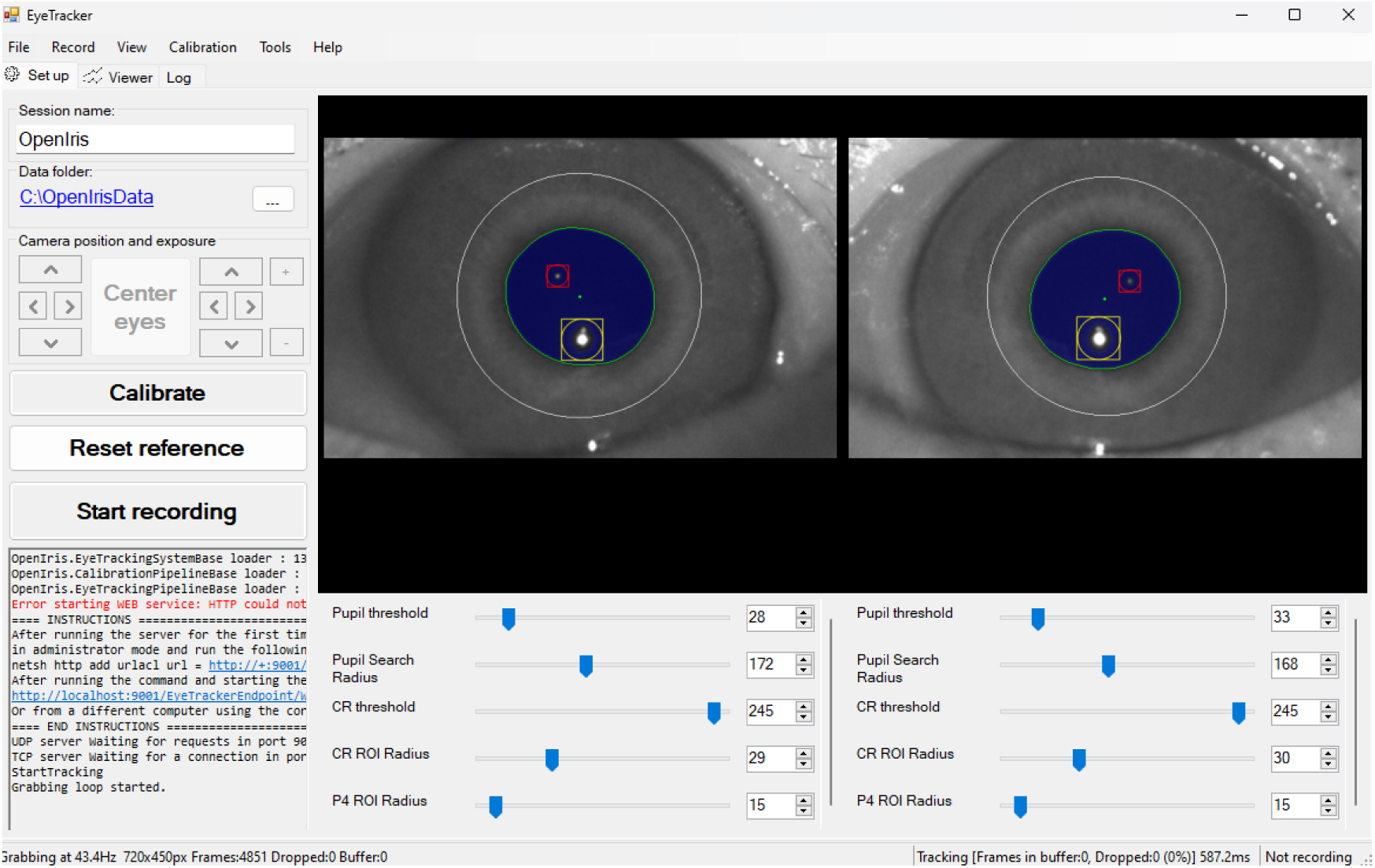
The OpenIris graphical user interface. The upper-right panel shows binocular images captured from an oblique angle. The tracking algorithm identifies three features in each eye: the corneal reflection (CR, yellow), the posterior lens reflection (P4, red), and the pupil contour (fitted ellipse, green). The white circle denotes the region where pixels below a threshold are assumed to belong to the pupil, and blue shading indicates pixels below that threshold. The small spot abutting the CR is the second Purkinje reflection (P2). The bottom panel provides interactive control of tracking parameters including thresholds, search regions, and feature detection settings. The left panel is used to configure data acquisition parameters—including camera and recording settings—and displays the system log.

There are two noteworthy limitations of the system that are common to DPI eye trackers. The first is its relatively narrow field of view. When the eye rotates by more than approximately 10^◦^ from rest, the fourth Purkinje image becomes blocked by the iris, limiting the range of trackable eye positions. The second limitation is inaccurate eye position estimates during periods of high acceleration, such as during saccades. When the globe accelerates sharply, the inertia of the lens and the elasticity of the suspensory ligaments conspire to produce an oscillatory motion of the lens, and thus P4. As a result, the DPI signal briefly decouples from the angular position of the globe during and immediately after a saccade (Deubel and Bridgeman 1995a).

Nevertheless, the OpenIrisDPI eye tracking system is particularly useful for studies for which small changes in eye position between saccades have a large effect on the dependent variable(s) of interest. These include studies of intersaccadic drift on perception, binocular coordination, and the neurophysiological influence of top-down visual signals, which can be difficult to dissociate from bottom-up signals in the absence of high-quality eye tracking. As described in the Results, we used the system to measure the spatial properties of receptive fields of neurons in the macaque lateral geniculate nucleus (LGN). This application leverages the exquisite sensitivity of the tracker to fixational eye movements, which inflate estimates receptive field size unless compensated.

While this system was developed for use with nonhuman primates, adapting the system for human use is possible with minimal difficulty, although additional care will be needed to limit risks from infrared exposure (see Discussion).

### 1.1. DPI eye tracking for visual neuroscience

Highly precise eye tracking systems are essential for visual neuroscience, where accurate knowledge of the retinal image is critical to characterize neural responses (D. M. Snodderly 2016). Neurons in the visual system respond to specific patterns of light falling on the retina (Hubel and Wiesel 1961; Hubel and Wiesel 1962; Levitt et al. 1994). The region of the retina that contributes to a given neuron’s response (called the receptive field, or RF) samples different portions of the visual world as the eye moves (Gur, Beylin, et al. 1997; Gur and D. M. Snodderly 1987; Xiao et al. 2024). The RFs of some neurons subtend just fractions of a degree, which is substantially smaller than both the amplitude of many eye movements and the noise in some eye trackers.

Experimentalists, therefore, often attempt to minimize eye movements during studies of visual processing, typically by training research animals to fixate during stimulus presentations. However, residual eye movements during fixation are inevitable and modulate neural activity throughout the visual hierarchy (e.g. Bair and O’Keefe 1998; Khademi et al. 2020; Leopold and Logothetis 1998; D. M. Snodderly et al. 2001). Compensating for the effects of these movements on the retinal image, either online (e.g. Gur and D. M. Snodderly 1987; D. Snodderly and Gur 1995), or in post-hoc analysis (e.g. Gur and D. M. Snodderly 2006; Livingstone and Tsao 1999; McFarland et al. 2014; Read and Cumming 2003; Tang et al. 2007), can improve measurements of neuronal responsiveness, stochasticity, and selectivity.

To demonstrate that OpenIrisDPI is suitable for such high-precision applications, we used the system to study the light response properties of neurons in the macaque LGN. These neurons were recorded as part of an ongoing study of saccade-related activity modulations which will be described in a separate report. In this report, we use these data to validate OpenIrisDPI for receptive field estimation. We show that DPI-based eye tracking improves the signal-to-noise ratio and spatial resolution of spike-triggered averages and outperforms P-CR-based eye tracking in fixating monkeys. These results highlight the practical impact of high-precision eye tracking for visual neuroscience and demonstrate that compensating for variability introduced by eye movements can lead to more accurate characterization of neural responses in the early visual system of macaque monkeys, even during nominal fixation.

## 2. Materials and methods

### 2.1. OpenIrisDPI algorithm

The OpenIrisDPI tracking algorithm processes video input sequentially: first fitting the pupil boundary, then identifying the CR, and finally localizing P4. Pseudocode for the algorithm is given in Appendix B.

The tracking algorithm is implemented as a plugin for OpenIris and utilizes the OpenCV computer vision library through its C# wrapper, Emgu CV. OpenCV is highly optimized for CPU-based parallelism, and enables OpenIrisDPI to process video at high rates. All data presented in this publication were processed in real time from two temporally synchronized video streams, each imaging a single eye at 500 Hz. This frame rate was achieved by reducing the region of interest from 1440×1080p to 720×450p. Although data were acquired binocularly, data from the left eye only were analyzed. The computer running OpenIrisDPI consisted of consumer-grade components, namely a 12th generation Intel Core i9-12900K CPU, 16 GB of DDR4 RAM, and a 512 GB SSD.

### 2.2. Eye tracking hardware and safety

The optical hardware configuration consists of two CMOS digital cameras (FLIR BFS-U3-16S2M-CS) equipped with 100 mm macro lenses (Laowa). These cameras were mounted below the subject’s line of sight (35°). Each camera was positioned to image a single eye at a distance of 57 cm. Infrared illumination was provided by a focused LED source (Thorlabs M940L3) positioned above the cameras (at an oblique angle of 25°) to optimize the visibility of the CR and P4 from both eyes. In this configuration, P4 was visible for eye positions up to 10° from central position in any direction before becoming obscured by the iris.

To ensure the safety of the infrared illuminator, we measured the irradiance of the focused infrared beam in front of the eyes to be 49 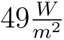 using a Thor-labs PM100D optical power meter with S121C sensor. This value is below the limit set by the IEC for indefinite corneal and lens exposure (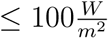, *Photo-biological safety of lamps and lamp systems* 2006). We also measured the peak irradiance, which was near the monkey’s nose to ensure even illumination of the two eyes, and found it to be 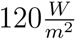. We further calculated the weighted retinal exposure at the irradiance peak to be 5363.15 W/m²/sr, which is more than a log unit below the safety limit for retinal burn set by the IEC. For further details on how these values were calculated, see the OpenIrisDPI wiki (https://github.com/ryan-ressmeyer/OpenIrisDPI/wiki/IR-Safety).

### 2.3. Model eye experiments

An artificial eye (Ocular Instruments OEMI-7) was used to benchmark measurement error. The model eye was attached to a stepper motor (STEPPERONLINE Nema 17) that was connected to a driver (STEPPERONLINE DM542T) that moved in 0.54-arcminute steps. The accuracy of these steps was validated by attaching a laser pointer to the motor and measuring the displacement of the beam over a distance of 43 feet. The artificial eye and stepper motor were placed on a platform at the approximate location of the monkey’s eyes during experiments. To simulate standard use, no effort was made to reduce vibrations of the eye or cameras. A third reflection, not present in real eyes, was visible in the model eye and ignored by the tracking algorithm. During experiments, the model eye was rotated through a series of five equally spaced steps from 2.16’ to 10.8’, with 2 s pauses at each step. This series was followed by 10 minutes of stationarity to quantify measurement noise with a static eye.

### 2.4. In vivo preparation and eye tracker calibration

All protocols conformed to the guidelines provided by the US National Institutes of Health and the University of Washington Animal Care and Use Committee. Data were collected from two adult rhesus macaques (Macaca mulatta), one male (Monkey O) and one female (Monkey C). Each monkey was surgically implanted with a titanium headpost and a recording chamber (Crist Instruments) stereotaxically positioned over the LGN. During experiments, the monkeys sat in a primate chair (Crist Instruments) 61 cm from a rear projected digital display (VPixx PROPixx, 51 cm width, 1280×720p resolution) in a dark room while their eyes were tracked using OpenIrisDPI. Monkeys were trained to sit and fixate projected black dots (0.2 *×* 0.2^◦^) for juice rewards. Data from 11 recording sessions (7 monkey O, 4 Monkey C) were analyzed for this study.

The experiments were conducted in two phases: calibration and fixation. During calibration, saccade targets (0.2 *×* 0.2^◦^) were presented in random order on a uniformly spaced 7 *×* 7 grid spanning 12 *×* 9^◦^ for Monkey O or in a 5 *×* 5 grid spanning 4 *×* 4^◦^ for Monkey C. This difference between monkeys met the requirements of an unrelated study and does not affect the conclusions of the this report. The monkey’s eye position was communicated from the OpenIrisDPI computer to the experimental control computer using a pair of manually-calibrated analog voltage signals generated by a USB digital-to-analog module (ACCESIO USB-AO16-8E). The analog signals were conditioned with a custom lowpass 8-pole Bessel filter (180 Hz cutoff) before re-digitization. A target was considered fixated if this signal indicated the eye position was within 1.4 *×* 1.4^◦^ of its displayed position for a duration of 600 ms. The target was then extinguished, and the monkey received a juice reward paired with a tone. The analog eye position signal was used for online experimental control only. All analyses presented in this paper were performed using the digital signals saved by OpenIris, which were digitally synchronized to event times in offline analysis.

Raw DPI and P-CR signals for the left eye were computed by taking the difference between the estimated centers of the CR and P4 (for DPI), or the CR and pupil (for P-CR). Data from the calibration task were processed as follows. First, the timestamps of the eye position signals were aligned to the times of task events (e.g. stimulus appearance) as described in the OpenIrisDPI wiki (https://github.com/ryan-ressmeyer/OpenIrisDPI/wiki/Synchronizing-OpenIris-Data). Then, the median DPI and P-CR pixel positions during the 100 ms prior to target offset were calculated. Outlier fixations were removed using a distance-based criterion, which was applied independently to the cluster of fixations for each target location. For each fixation within a cluster, we calculated its average distance to all other fixations within that same cluster. This process yielded a set of average-distance scores, one for each fixation. Any fixation for which the average distance score was more than the median+3*×*IQR of all scores was discarded. The remaining points were then regressed onto the positions of the displayed target positions using a quadratic model (Kimmel et al. 2012). The quality of the fits were visually inspected for accuracy. Where appropriate, calibrated signals are reported in units of visual angle using a small angle approximation.

### 2.5. Neurophysiological experiments

After calibration, the fixation task began. During the task, a black dot was presented continuously at a set location, typically the center of the screen, to ensure that the recorded LGN neurons could be stimulated by the display. The monkey was rewarded at semi-random intervals (range 400–800 ms) for maintaining fixation on this dot. While the monkey fixated, a 120 *×* 120-element stimulus grid subtending 17.3*×*17.3^◦^ was displayed over the receptive fields of recorded neurons. The RGB values of each element were updated at 60 Hz using independent draws from a Gaussian distribution with mean equal to the background. The task continued until 12–18 minutes of stimulus frames had been presented.

We recorded extracellular action potentials from LGN neurons using high-density silicon probes (Neuropixels NHP 1.0 long). The probes were held using a custom adapter that maintained alignment between the probe shank, a guide tube (25 mm-long 20G hypodermic needle), and microdrive assembly (Narishige). Before experiments began, the guide tube was lowered through the dura mater using a thumb screw. The probe was then advanced using a hydraulic microdrive until it penetrated the LGN. Neural activity was verified as originating from the LGN using a hand-held flickering red light. The action potential band (sampled at 30 kHz) and local field potential band (sampled at 2.5 kHz) were recorded using OpenEphys (Siegle et al. 2017). Action potentials were sorted offline using Kilosort2.5 (Pachitariu et al. 2024).

### 2.6. Inter-saccadic drift analyses

Eye position analyses were restricted to epochs when the eye position was within 1.5° of the fixation point for at least 500 ms. Saccades during these epochs were identified using a standard algorithm (Nyström and Holmqvist 2010), and intersaccadic epochs were identified as periods between saccades that were ≥ 50 ms. Intersaccadic epochs were ignored if the maximum dis-placement exceeded 1° in a 200 ms window.

The power spectral density (PSD) of horizontal and vertical eye position during fixations was estimated separately for each experimental session using Bartlett’s method as implemented by the Python library, Scipy. The square root of the PSD was computed to yield the amplitude spectrum. Diffusion constants were estimated by computing the mean squared displacement (MSD) of eye position for time lags of 2 to 200 ms during fixations, then taking the slope of the best-fit line and dividing it by 4 (A. M. Clark et al. 2022; Herrmann et al. 2017; Intoy and Rucci 2020).

### 2.7. Spike-triggered analysis

We analyzed neural responses to the white noise stimuli using spike-triggered averaging. Eight stimulus frames preceding each spike were reconstructed offline and averaged across spikes (Chichilnisky 2001). This operation yielded the spike-triggered average (STA) as a four-dimensional tensor, STA_l,c,y,x_, where the indices correspond to time lag (l), color channel (c), vertical (y), and horizontal (x) coordinates.

To determine which STAs contained reliable signatures of visual responsiveness, we calculated a signal-to-noise ratio (SNR) for each STA. First, we computed the magnitude of the STA at each spatiotemporal pixel, I_l,y,x_, by taking the L2-norm across the three color channels (Eq. (1)). The signal was defined as the maximum magnitude observed across all time lags and spatial positions. The noise was estimated as the median magnitude within the first time lag (Eq. (2)). This choice of noise reference is justified by the minimum response latencies of macaque LGN neurons (≈16 ms), which ensures the first stimulus frame is non-causal (Maunsell et al. 1999). Units—the clusters found by Kilosort—that produced an STA with SNR below 5 were excluded from further analyses.

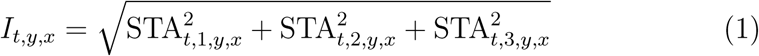

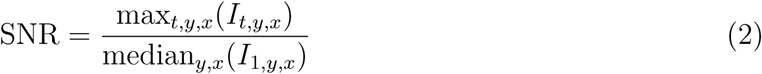

To account for fixational eye movements, we reconstructed a gaze-contingent version of the stimulus offline to approximate the image on the retina. First, we processed the DPI and P-CR eye-tracking signals using a zero-phase 8th order Chebyshev filter (30 Hz cutoff) and interpolated them to find the gaze position at the midpoint of each stimulus frame. Each frame used for the STA calculation was then spatially shifted to align with the estimated gaze position at that time.

For STAs computed with and without gaze correction, we fit a 2D Gaussian (Scipy curve fit) to the single frame that had the highest value of I_l,y,x_. STAs were excluded if the fit failed to converge or yielded implausibly large radii (> 20^′^). The final RF radius was calculated as the geometric mean of the standard deviations along the fitted major and minor axes.

## 3. Results

### 3.1. Innovations of OpenIrisDPI

Significant contributions of OpenIrisDPI are (1) a CPU-optimized algorithm for simultaneously tracking the pupil, CR, and P4 in real time, (2) documentation of hardware configurations (Fig. 3), including an oblique configuration that is novel for DPI tracking, and (3) integration with the OpenIris eye tracking framework for ease of use and extensibility.

**Figure 3:**
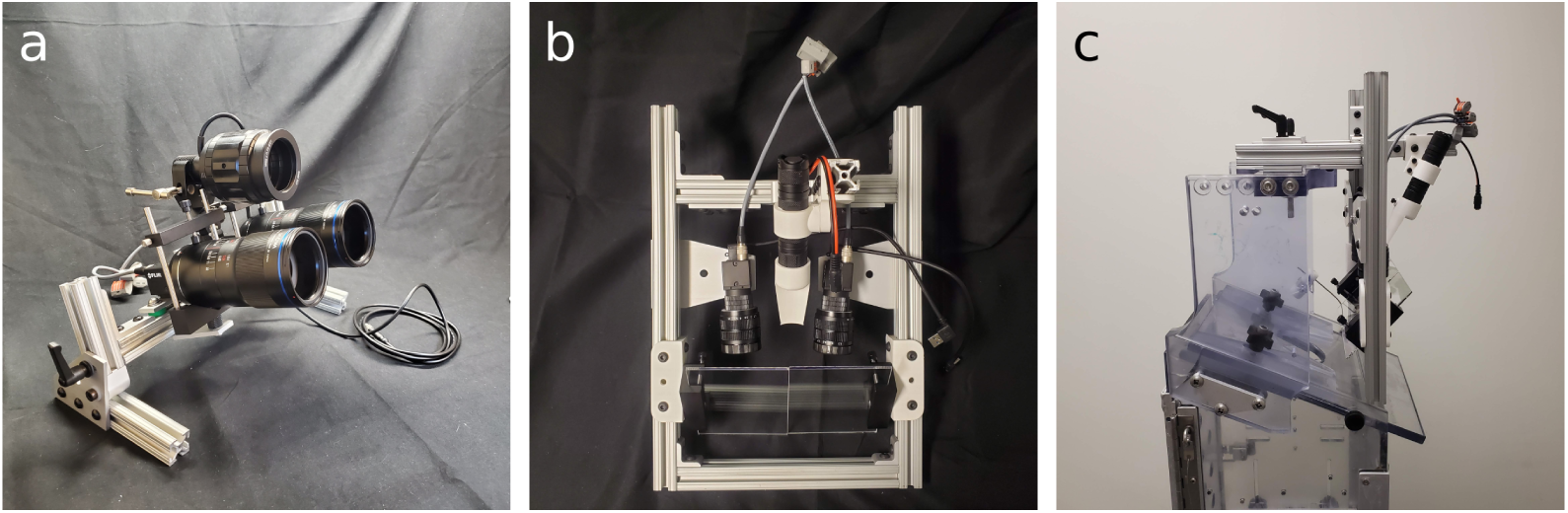
Hardware configurations for DPI eye tracking. a) An oblique imaging system positioned below the subject’s line of sight that does not require a hot mirror. b) A conventional, paraxial imaging system that uses hot mirrors to align the optical axes of the cameras and the eyes. c) The paraxial imaging system mounted to a Crist Instruments primate chair.

Most previously described DPI eye trackers use a hot mirror to align the optical axes of the eye and camera (Abdulin et al. 2019; Crane and Steele 1985; Wu et al. 2022). This approach minimizes signal non-linearity while still allowing the participant to view experimentally controlled stimuli. However, a hot mirror also adds complexity by filtering the light between the display and the subject. We found that robust DPI eye tracking is achievable from an oblique angle provided that the infrared illuminator is placed at a shallower angle than the imaging cameras. This configuration centers P4 in the pupil for neutral gaze, which maximizes the range of eye positions over which P4 is visible. Placing the imaging system at an oblique angle eliminates the need for a hot mirror and allows a DPI eye tracker to be a drop-in replacement for many commercial systems. All data presented in this study were collected using the oblique imaging configuration.

A second innovation of OpenIrisDPI is its CPU-optimized, real-time tracking algorithm. Digital DPI eye tracking systems described previously either process video data post-hoc (Tabernero and Artal 2014) or rely on GPUs to track the Purkinje reflections in real-time (Wu et al. 2022). By optimizing the OpenIrisDPI algorithm for the CPU, we aimed to make DPI eye tracking portable and cost-effective, with minimal performance trade-offs. To achieve this goal, we compared the radially symmetric center algorithm for reflection tracking (Parthasarathy 2012) implemented by Wu et al. 2022 to a threshold-rectified centroid algorithm (Ares and Arines 2004). The latter was faster and more accurate for reasonable thresholds in simulations (Fig. B.1). In addition, we simplified the algorithm for locating P4 relative to the algorithm presented by Wu et al. 2022, which involves an expensive template matching operation. Excluding the CR, P4 is typically the brightest object in the pupil. We therefore locate P4 by identifying the maximum amplitude pixel within the pupil border, excluding a small region surrounding the CR. The result of these optimizations is a lightweight algorithm that is capable of binocularly tracking the Purkinje reflections and the pupil simultaneously at 500 Hz using consumer-grade hardware. This supports both high-resolution gaze tracking and simultaneous pupillography.

We characterized the processing time of the algorithm, as well as the end-to-end latency between exposure offset and the analog output. Under our current system, the time to process one frame from each camera is 1.1±0.1 ms (median ± IQR). Occasionally, however, the processing time was appreciably longer, up to a maximum of 50 ms. These infrequent (2% of frames ≥ 10 ms) processing delays are likely due to CPU preemption by background operating system processes. The real-time analog output signal inherited these delays and imposed an additional 3–4 ms delay, which may limit the use of this signal for gaze-contingent applications. Importantly, every frame from both cameras was timestamped, stored to disk, and registered to the times of experimental events. This allowed us to reconstruct the eye position signals with high temporal precision offline irrespective of occasional processing delays (see Methods).

### 3.2. DPI signals exhibit sub-arcminute precision with a model eye

To assess the baseline performance of the OpenIrisDPI system in our experimental configuration, we used a model eye affixed to a stepper motor (Fig. 4a). The eye was driven through a series of 2.16–10.8’ steps for calibration (Fig. 4b), then held steady for 10 minutes to assess signal stability. During these 10 minutes, the DPI signal was less variable than the P-CR signal (Fig. 4c; P-CR RMS: 1.83’; DPI RMS: 0.27’). The amplitude spectrum of both signals was flat above 50 Hz and increased at lower f<u>req</u>uencies. The median amplitude of the P-CR signal (1.83 *×* 10^−3^deg / 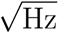) was similar to that obtained by Niehorster et al. 2021 using the SR EyeLink 1000 Plus, and the median amplitude of the DPI signal (2.11 *×* 10^−4^deg / 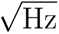) was substantially lower than any eye tracker reported in that study.

**Figure 4:**
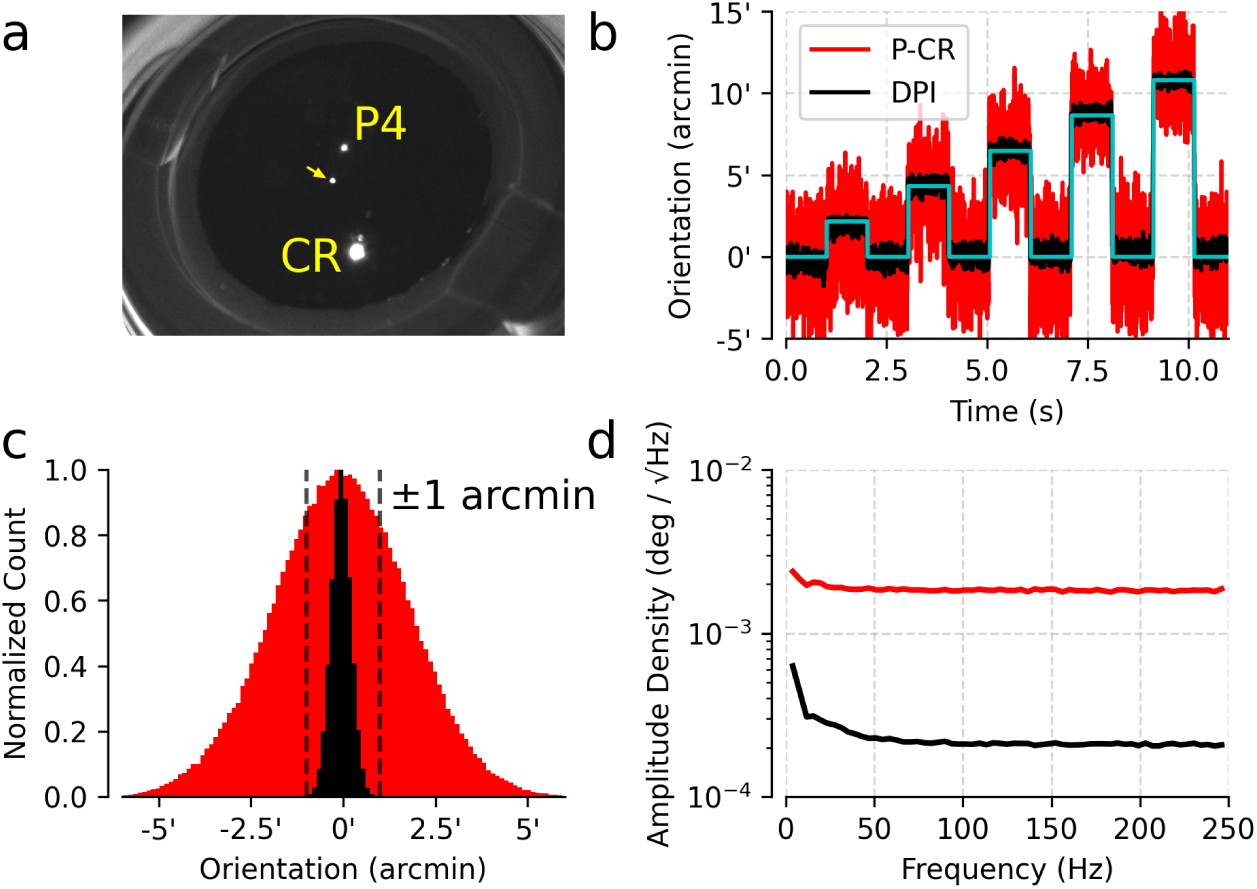
Tracking an artificial eye. a) Single frame from a video of the model eye. CR and P4 are labeled in yellow. A reflection from the model eye is absent from real eyes (arrow). b) P-CR (red) and DPI (black) signals in response to a series of steps. The cyan line indicates the requested rotation. c) Distribution of reported orientation while the model eye was stationary. d) Amplitude spectra of P-CR and DPI signals.

Both P-CR and DPI signals had higher amplitude at low than high frequencies (3.91 Hz amplitude minus median amplitude; DPI: 2.90*×*10^−4^ deg / 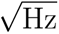; P-CR: 2.35 *×* 10^−4^ deg / 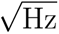). This increase in low frequency power was presumably caused by ambient mechanical oscillations present in the experimental setup.

### 3.3. Comparing DPI and P-CR signals in vivo

Model eyes, while commonly used for eye tracker validation, fail to capture many aspects of real eyes including pupillary decentration (Wyatt 1995), saccade-related iris kinematics (Nyström, I. Hooge, et al. 2013), lens wobble (Deubel and Bridgeman 1995a), and ocular tremor (Spauschus et al. 1999). Consequently, measures of precision obtained with model eyes are often overestimates (Wang et al. 2017). We therefore compared DPI and P-CR measurements from two rhesus macaques (see Methods). In addition to a small DC offset, the resultant eye traces (Fig. 5) displayed (1) greater frame-to-frame noise in the P-CR signal than in the DPI signal, and (2) an additional slow drift in the P-CR signal that the DPI signal lacked. The following analyses isolate and quantify these two effects.

**Figure 5:**
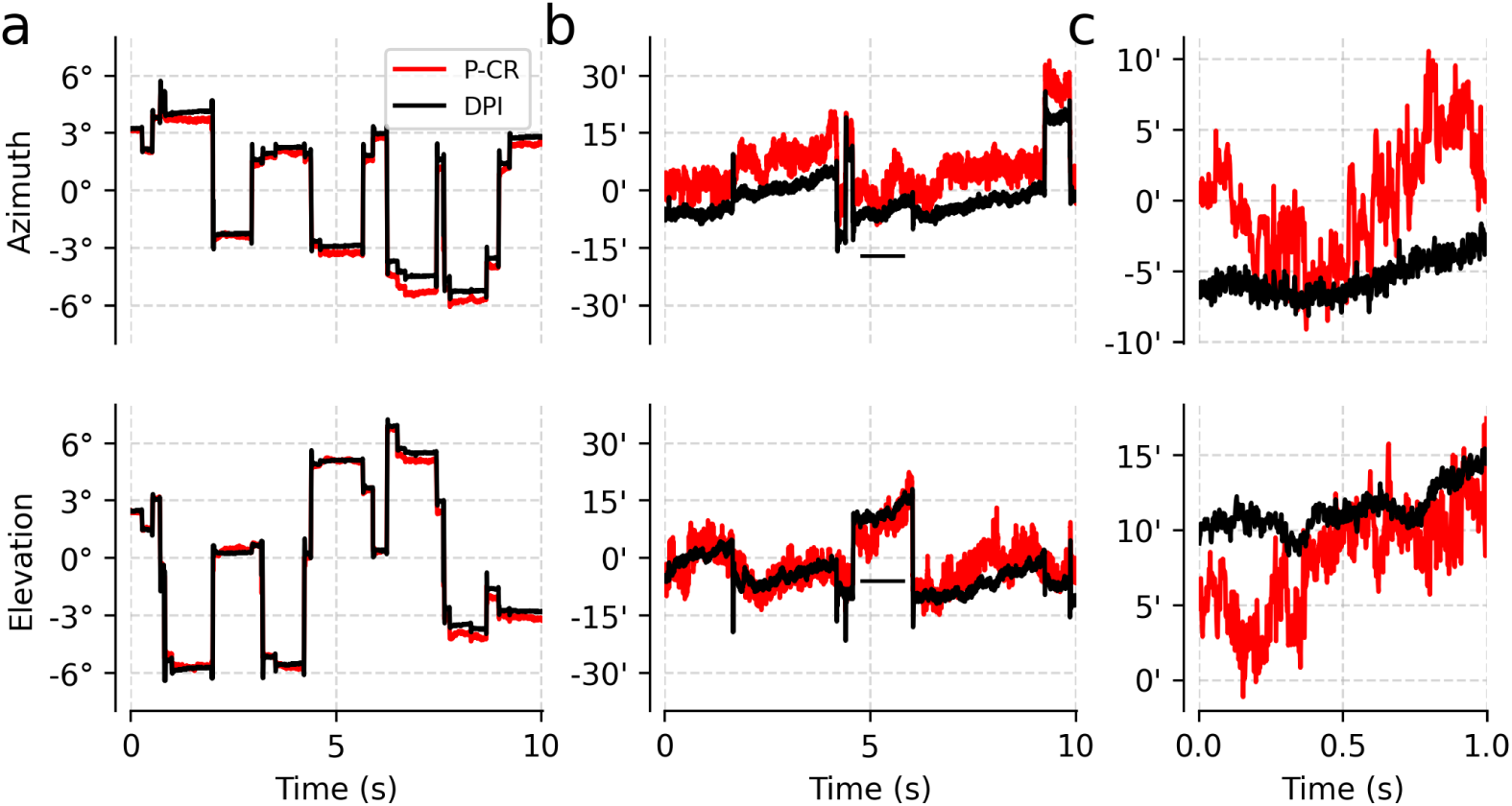
Example of simultaneously recorded DPI (black) and P-CR (red) measurements of eye azimuth (top) and elevation (bottom). a) Eye position signals during the calibration task. b) Eye position signals during the fixation task. Black bar indicates time window shown in (c). c) Eye position signals during an inter-saccadic period. Note difference in ordinate scales: the y-axis for (a) is in degrees, while for (b & c) it is in arcminutes.

#### 3.3.1. DPI signals exhibit less frame-to-frame noise than P-CR signals

To quantify frame-to-frame measurement noise of the DPI and P-CR signals, we calculated the amplitude of broadband noise between saccades. We assume that eye tracking signals, x[n], are the sum of a noiseless signal, s[n], and a temporally uncorrelated, zero-mean noise source, ɛ[n]. We are interested in estimating the standard deviation of 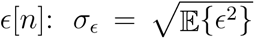. If we had access to ɛ[n] in isolation, we could use the RMS as an estimate. However, since we only have access to the combined signal, *x*[n], another method is needed. If s[n] is band-limited below a frequency cut-off, f_c_, then an estimate of σ_ɛ_can be obtained from the average spectral power above *f*_c_ (Schuster et al. 2019). We obtain this estimate using the formula:

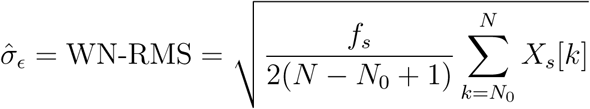

where X_s_[k] is a normalized, one-sided estimate of the PSD of x[n], N is the number of bins in the PSD, N_0_ is the first bin above f_c_, and f_s_ is the sampling frequency of the signal. We refer to this measure as the white noise RMS (WN-RMS) amplitude.

We calculated the WN-RMS amplitude of DPI and P-CR signals to quantify frame-to-frame noise during our electrophysiological experiments (n=11). The DPI signal exhibited a broad peak in spectral power around 100 Hz, which is consistent with prior reports of ocular tremor (Ko et al. 2016). Therefore, f_c_ was chosen to be 200 Hz to minimize contamination by this signal (Fig. 6a). Given this threshold, the WN-RMS amplitudes of the DPI signals (WN-RMS_dpi,az_ = 0.39^′^*±*0.06^′^, WN-RMS_dpi,el_ = 0.44^′^*±*0.07^′^) were ap-proximately a third of the amplitude of the P-CR signals (WN-RMS_p-cr,az_ = 1.55^′^ *±* 0.20^′^, WN-RMS_p-cr,el_ = 1.82^′^ *±* 0.26^′^) (Fig. 6b). These differences were statistically significant (p < 10^−3^, Wilcoxon signed-rank test). Furthermore, the noise amplitude of the azimuthal components of the DPI and P-CR signals were consistently lower than that of the elevational components (p < 10^−3^, Wilcoxon signed-rank test), which may be due to the oblique optical configuration. Finally, we observed a significant difference in P-CR WN-RMS amplitude between Monkey C and Monkey O (p < 10^−2^, Mann-Whitney U test on mean of WN-RMS_az_ and WN-RMS_el_), which may be due to differences in head position between the two animals. There was no statistically significant difference in DPI noise floors between the two monkeys (p = .16, Mann-Whitney U test). Thus, the data confirm that DPI signals are less susceptible to temporally uncorrelated measurement noise than P-CR signals when computed from identical video inputs, at least when using the OpenIrisDPI system under the conditions of this study.

**Figure 6:**
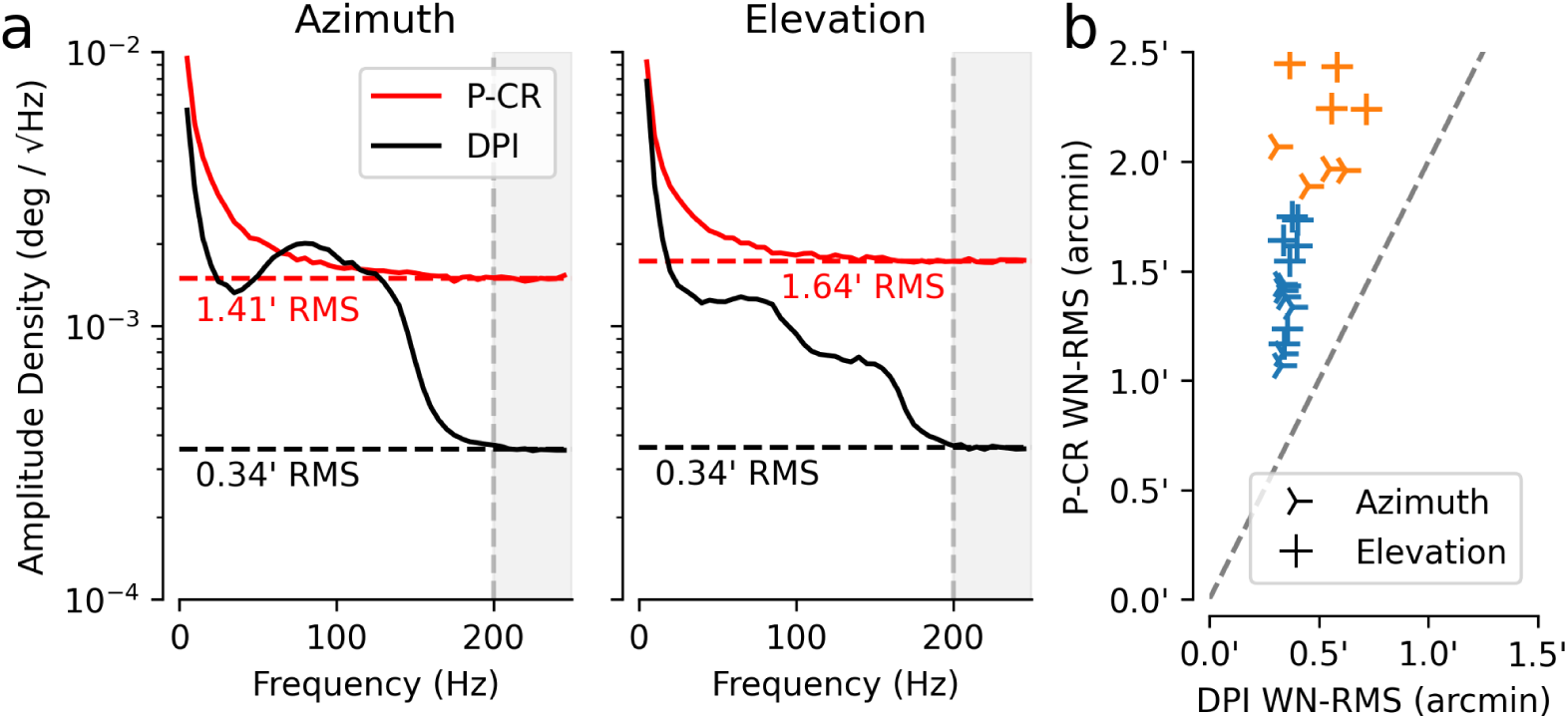
White noise amplitude during intersaccadic drift. a) Amplitude spectrum of DPI (black) and P-CR (red) signals for azimuthal (left) and elevational (right) components from an example session. Dashed lines indicate the mean spectral amplitude density above 200 Hz (shaded gray). b) WN-RMS value computed for DPI (abscissa) and P-CR (ordinate) signals. Azimuthal and elevational signal components are marked with three-pointed symbols and crosses, respectively. Blue and orange markers indicate data collected from Monkey O and C, respectively. The gray dashed line indicates *y* = 2*x*.

#### 3.3.2. P-CR signals report more ocular drift between saccades than DPI signals

In addition to having greater high-frequency noise, P-CR signals also drifted more than DPI signals do (Fig. 7a & b). One possibility is that this excess drift was due to biological factors such as pupil decentration (Wyatt 2010). To address this possibility, we once again assumed that measurement noise is additive and uncorrelated between successive frames, and that signals of biological origin vary slowly over time. Under these assumptions, at every moment in time the estimated eye position is the sum of a slow component (whose deviations accumulate over time, like Brownian motion) and measurement noise that does not accumulate over time. To isolate this slow component, we calculated the mean squared displacement (MSD) as a function of lag and compared its magnitude between DPI and P-CR signals (Fig. 7c).

**Figure 7:**
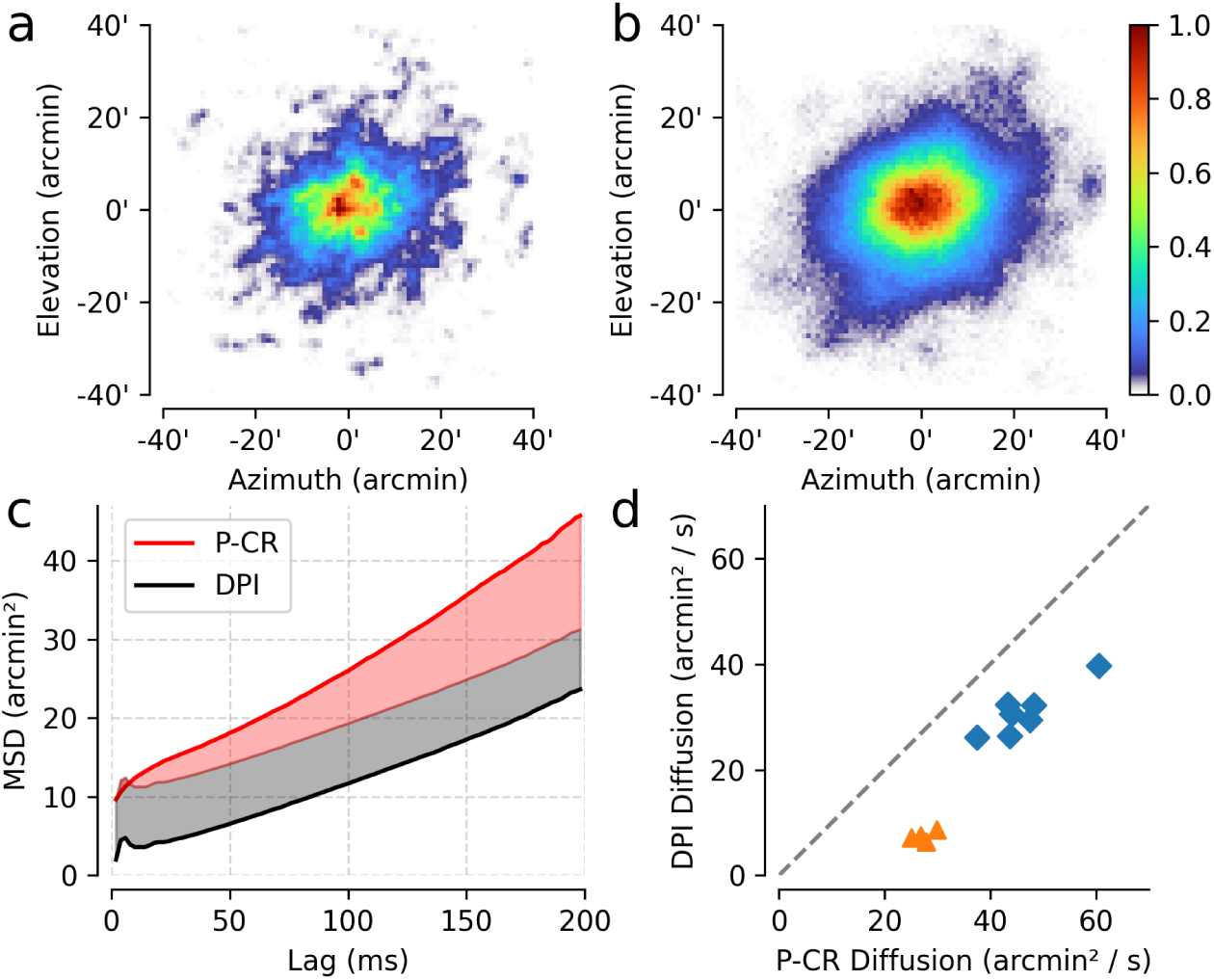
P-CR signals are more dispersed than DPI signals during intersaccadic drift. a) Normalized distribution of gaze direction during fixations measured using DPI. b) Same as in a, but using simultaneously collected P-CR data. c) Mean squared displacement (MSD) over various lags for DPI (black) and P-CR (red) signals calculated during intersaccadic drift. Temporally uncorrelated noise can cause a vertical offset between the two traces (shaded gray) but cannot produce the obsevered difference in slope (shaded red). d) Diffusion coefficients estimated using P-CR (abscissa) and DPI (ordinate). Blue squares indicate data from monkey O, orange triangles indicate data from monkey C.

MSD as a function of temporal lag is commonly used to estimate the parameters of a Brownian motion model of intersaccadic drift (e.g. Engbert and Kliegl 2004). Under this model, the slope of the line of best fit is four times the diffusion coefficient of the Brownian motion (Intoy and Rucci 2020). We presume that the motion of the eye is identical whether measured by P-CR or by DPI, so we expect that this slope will be the same for both techniques. Note that stationary white noise increases the MSD for all lags equally, and as a result this prediction holds even if the level of measurement noise differs between techniques. What we observed, however, is that the diffusion coefficient was 17.12 ± 1.09 arcmin²/s larger (p < 10^−4^, Wilcoxon signed-rank test) when calculated from P-CR data than from DPI data (Fig. 7d), indicating that the P-CR signal contains additional low-frequency drift that is distinct from its elevated white noise floor.

### 3.4. DPI eye tracking improves receptive field characterization in fixating macaques

We sought to demonstrate the benefits of using a DPI eye tracker for in vivo neurophysiology by analyzing responses to visual stimulation in the LGN of macaque monkeys. Neurons in the LGN receive direct input from the retina and have punctate receptive fields that range from several to tens of arcminutes in diameter (Croner and Kaplan 1995). We recorded from populations of LGN neurons while the macaques were tasked with fixating a stationary point (Fig. 8a). During periods of stable fixation, a white noise stimulus was displayed over the recorded population’s receptive fields. In offline analysis, we attempted to compensate for small eye movements made during fixation using either the DPI or P-CR signals recorded by OpenIrisDPI (Fig. 8b).

**Figure 8:**
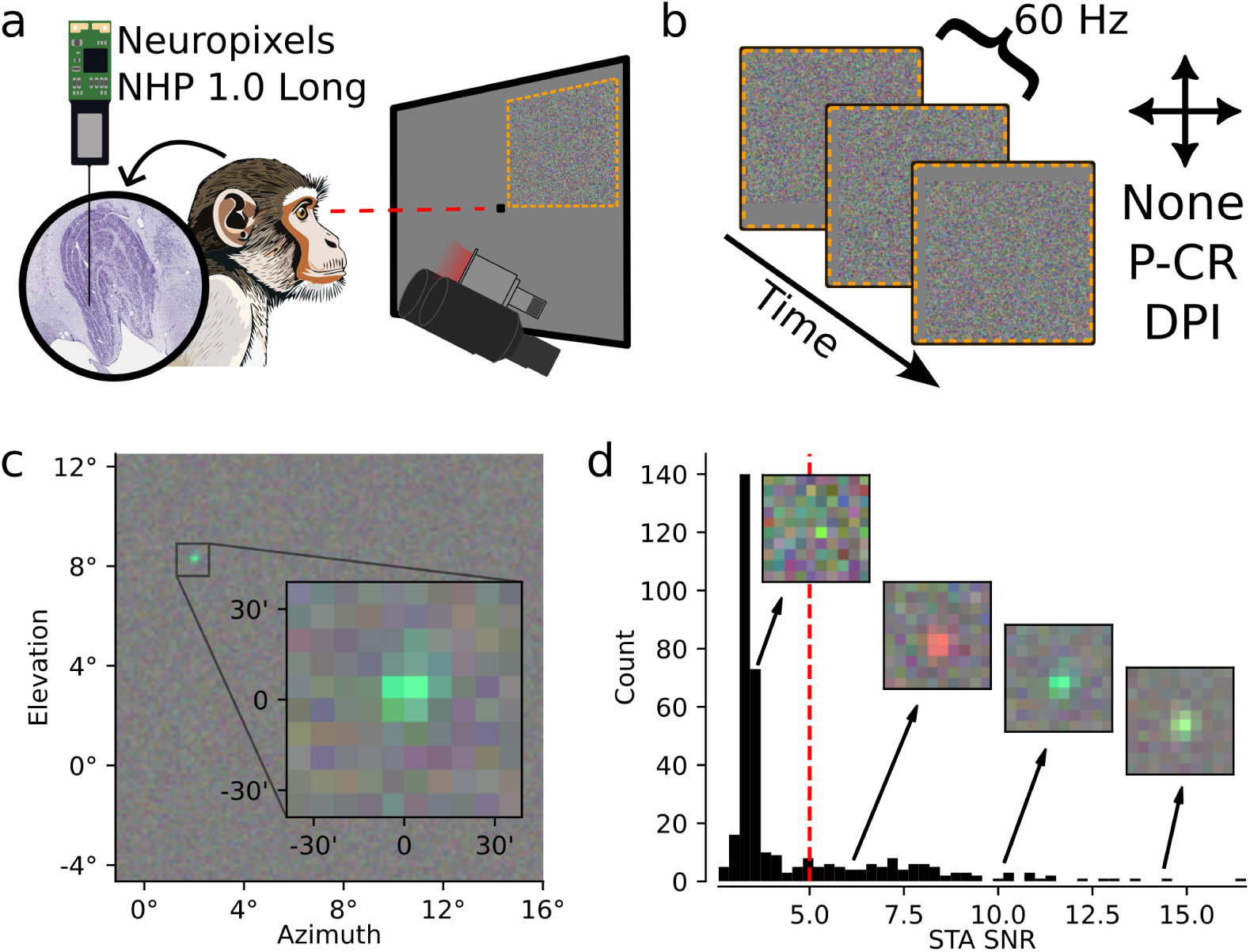
Electrophysiological experimental design and analysis. a) A 45 mm-long Neuropixels NHP 1.0 probe was inserted into the LGN of a macaque fixating a centrally projected dot (gaze drawn in red). Gaze direction was monitored using OpenIrisDPI. During periods of fixation, a white noise stimulus was displayed over the receptive fields of the LGN neurons recorded by the probe (dashed orange square). b) During analysis, the average stimulus preceding the spikes of each unit was calculated. Each frame of the stimulus was then shifted to account for the effects of fixational eye movements on the retinal movie, and the analysis was repeated. c) The peak lag of a representative spike-triggered average (STA) of a presumptive parvocellular neuron without correction for residual eye movements. Inset shows the STA cropped around the RF. d) Histogram of signal-to-noise ratio (SNR; see Eq. (2)) for each unit in a representative recording. Insets show cropped STAs, and arrows point to the corresponding SNRs. Only STAs with SNR exceeding a threshold (red dashed line) were included in subsequent analyses. Inset in (a) reproduced from brainmaps.org (Mikula et al. 2007).

Reverse correlating the recorded spikes with these compensated stimuli showed that DPI signals greatly improved estimates of neuronal tuning compared to either no compensation—that is, ignoring eye movements—or compensation using P-CR signals. We first computed the spike-triggered average (STA) of every unit identified by Kilosort (average 405 units per recording; range 176–638 units per recording), and computed the SNR of the resulting STAs (see Eq. 2). Across experimental sessions, an average of 68 STAs (range: 33–133) had SNR ≥ 5 without gaze compensation (Fig. 8c & d). We then computed gaze-compensated STAs by translating each stimulus frame by whole-element amounts to account for the gaze positions reported by either DPI or P-CR signals. This revealed that both DPI and P-CR signals improved RF estimates, and that DPI-based compensation significantly outperformed compensation using P-CR signals (Fig. 9). The SNR of STAs relative to uncorrected data improved by 87.4% ± 10.0% for DPI correction versus just 29.0% ± 9.2% for P-CR (p < 10^−3^, Wilcoxon signed-rank test). Similarly, DPI correction produced significantly smaller receptive fields in 10 of 11 datasets, with RF radii contracting by 1.66’ ± 0.38’ on average for DPI versus 0.56’ ± 0.27’ for P-CR correction (p < 5 *×* 10^−3^, Wilcoxon signed-rank test). These analyses confirmed that OpenIrisDPI can be used to compensate for fixational eye movements and that DPI signals are superior to P-CR signals for this application.

**Figure 9:**
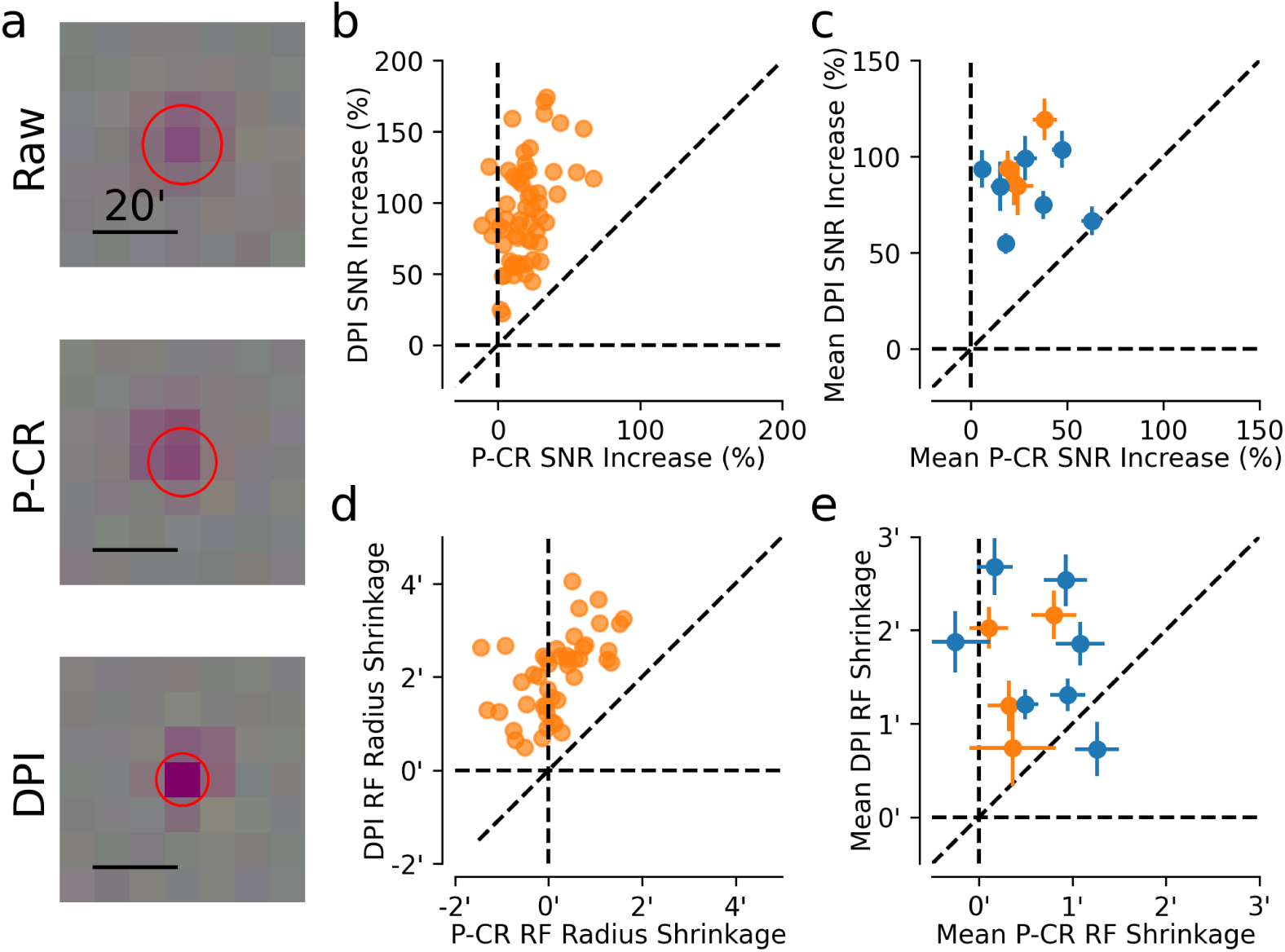
Comparison of DPI- and P-CR-compensated receptive field measurements of LGN neurons in fixating macaques. a) STA of an example unit with no compensation (top), P-CR compensation (center), and DPI compensation (bottom). Red circle indicates one *σ* of a Gaussian fit. Scale bar is 20 arcminutes. b) STA SNR improvement relative to baseline for all units with valid receptive fields in an exemplar recording using P-CR (abcissa) and DPI (ordinate) signals. c) Mean STA SNR improvement for all datasets. Error bars represent *±* 1.96 SEM. d) Reduction of STA radius compared to baseline for all units shown in b) using P-CR (abcissa) and DPI (ordinate) correction. e) Mean reduction of STA RF radius for each dataset. Error bars same as in (c). Color indicates animal identifier as in Fig. 7.

## 4. Discussion

We have described a new open-source eye tracking system, OpenIrisDPI, with the goal of increasing access to DPI eye tracking for visual neuroscience. DPI eye tracking is a non-invasive and highly precise technique that faces limited adoption due to a lack of convenient, cost-effective, and widely available systems. OpenIrisDPI enables researchers to construct custom DPI systems using inexpensive, easily obtained consumer electronics. The software required to operate it is free and includes a graphical user interface alongside common features of commercial eye tracking systems (Sadeghi et al. 2024). As a result, OpenIrisDPI significantly reduces the barriers to the adoption of DPI eye tracking.

### 4.1. Validation of OpenIrisDPI’s performance

To validate the usefulness of OpenIrisDPI for high-precision applications, we tested its performance using an artificial eye. The noise amplitude of the P-CR signal was equivalent to that of the Eyelink 1000 Plus reported by Niehorster et al. 2021, and the DPI signal was an order of magnitude less noisy. We note that our DPI eye tracker was approximately four times noisier than the one developed by Wu et al. 2022. This may have been due to the fact that we used lower magnification to track the pupil, which was necessary for the direct comparison of DPI and P-CR signals.

We then compared DPI and P-CR signals recorded from fixating macaque monkeys by isolating two distinct components of the eye position signal between saccades: temporally uncorrelated (white) noise and slowly varying drift. The white noise signal amplitude of P-CR signals was three times greater than those of simultaneously recorded DPI signals. Since both signals were calculated from identical video inputs, we can exclude hardware differences as causative. We conclude that DPI tracking is less prone to measurement noise than P-CR tracking when applied to living eyes, but we cannot rule out the possibility that improvements to the pupil fitting algorithm might narrow this gap.

Our analysis of slow variations in eye position showed that P-CR signals drift more than DPI signals. These results are consistent with prior reports comparing simultaneously collected P-CR and scleral search coil signals (Kimmel et al. 2012; McCamy et al. 2015). The drift we observed cannot be attributed to sensor noise and likely reflects a biological process. Since both DPI and P-CR signals use the CR, the increased fixational instability we observed must result from drift of the pupil center relative to P4. This conclusion is consistent with previous studies in which varying background illumination caused P-CR eye trackers to report artifactual deviations in eye position (Drewes et al. 2014; Kimmel et al. 2012; Wildenmann and Schaeffel 2013; Wyatt 2010). Our results extend this understanding by demonstrating that ongoing pupillary dynamics cause increased drift even when background luminance is constant. Some studies have used P-CR eye trackers to estimate the parameters of intersaccadic drift (Di Stasi et al. 2013; Engbert and Mergenthaler 2006; Herrmann et al. 2017). It may be necessary to revisit the conclusions of such studies in light of these results (also see Chen and Hafed 2013).

It is worth noting that our analyses focused on low-amplitude eye movements between saccades because DPI is ill-suited for measuring saccade kinematics. The rapid acceleration and deceleration of the globe during saccades cause the lens to oscillate, which manifests as ringing in the DPI trace at the beginning and end of saccades (Deubel and Bridgeman 1995a). This ringing is not representative of the angular position of the globe. Nevertheless, lens wobble is expected to transiently shift the image on the retina (Tabernero and Artal 2014), and has been shown to alter perceptual judgments accordingly (Deubel and Bridgeman 1995b). It is therefore a non-negligible aspect of vision, yet it remains relatively underexplored. The widespread adoption of precise DPI systems, facilitated by open-source platforms like OpenIrisDPI, may foster new lines of inquiry into how these intrasaccadic lens dynamics contribute to visual processing.

### 4.2. A DPI eye tracker for visual neuroscience

Our neurophysiological recordings from the macaque LGN demonstrate the value of OpenIrisDPI for visual neuroscience research. Awake macaques are a gold-standard animal model for studying primate vision. However, ocular fixation is imperfect even in well-trained macaques, and residual eye movements can affect measurements of neuronal tuning to visual stimuli. We show that OpenIrisDPI can be used to compensate for these fixational eye movements, as evidenced by the increase in signal-to-noise ratios and contraction of receptive field estimates.

Previous studies of visual receptive fields have attempted to compensate for the effects of fixational eye movements on receptive field estimates using other eye tracking systems with varying degrees of success. Tang et al. 2007 used search coils sutured to the sclera. They reported a reduction in RF widths of 4 arcminutes in V1 and the LGN, which is consistent with our results. However, Read and Cumming 2003 used a similar technique and discovered significant inaccuracies in the coil system that counteracted any beneficial effects. A similar result was reported by McFarland et al. 2014, who inferred the eye’s position using neural activity and found it to be inconsistent with scleral search coils signals. Using search coils to compensate for fixational eye movements is therefore possible but unreliable, which is concerning given the invasive nature of this technique. OpenIrisDPI, in contrast, is both noninvasive and straightforward to implement, as proven by its successful deployment in five laboratories at the time of publication.

OpenIrisDPI may also enable new paradigms for studying neural function in awake animals. Yates et al. 2023 showed that DPI eye tracking can be used to study foveal receptive fields in V1 of freely-viewing marmosets. Studies that build on this fixation-free methodology are poised to shed light on phenomena at the intersection of natural oculomotor behavior and visual processing, including saccadic suppression (Matin 1974) and oculomotor-related visual illusions (Otero-Millan et al. 2012; Troncoso et al. 2008). Furthermore, cognitive processes, like visual attention, impact the statistics of fixational eye movements (Engbert and Kliegl 2003; Hafed and J. J. Clark 2002; Intoy and Rucci 2020) and modulate neural activity in early visual areas (Reynolds et al. 2000; Roelfsema et al. 1998). High-precision eye tracking can be used to disambiguate the feed-forward effects of eye movements from modulatory feedback and thereby benefit the study of cognitive processes.

OpenIrisDPI may also be a useful tool for studying binocular coordination and vergence eye movements. Recent work by Jaschinski 2016 and by I. T. Hooge, Hessels, et al. 2019 demonstrated that artifacts caused by pupil-decentration are particularly troublesome when measuring vergence eye movements using pupil-based video eye trackers. DPI eye trackers overcome this limitation by being insensitive to unwanted pupillary motion. OpenIrisDPI’s ability to track both eyes simultaneously and to serve as a convenient, cost-effective replacement for existing pupil-based systems make it an appealing option for studying binocular eye movements with high precision.

The DPI tracker described in this report can be used to track the eyes human participants, but care must be taken to maintain safe levels of ocular irradiation. While the safety measurements we performed ensure that our eye tracking system was in compliance with widely used international safety standards (*Photobiological safety of lamps and lamp systems* 2006), the components described in this report are capable of over-irradiating the eye. Users are urged to measure the light power at the subject’s eye with an infrared power sensor prior to extended use in humans.

As an open-source project, OpenIrisDPI is designed to be modified and extended by its users. Several limitations of the current implementation present opportunities for future development. For example, the system’s narrow tracking range (∼ 10^◦^) could be expanded by incorporating multiple illuminators and an algorithm to switch between them or track their Purkinje reflections simultaneously. An alternative approach could pair a single illuminator with multiple cameras, ensuring that the CR and P4 reflections are resolvable in at least one camera across a wider range of gaze angles. Furthermore, OpenIrisDPI’s support for both P-CR and DPI tracking creates the potential for a hybrid system that dynamically switches between methods, leveraging DPI for high-precision fixational measurements and P-CR for lower-precision estimates of larger gaze angles. Other opportunities for development include overcoming the need for stable head fixation through a head-mounted imaging system, and mitigating tracking inaccuracies during saccades by developing more sophisticated models of the inertial properties of the lens.

Further refinements could focus on optimizing the imaging hardware itself. We selected an illuminator that produces minimally visible, 940 nm-light, but our camera’s quantum efficiency is less than 10% at this wavelength. Sourcing a camera with greater infrared sensitivity or using a shorter-wavelength illuminator could improve the signal-to-noise ratio and reduce risks associated with ocular irradiation. However, to our knowledge, no consumer-grade cameras with substantially greater IR sensitivity also offer the high framerates needed for 500 Hz eye tracking. By making the software and hardware specifications publicly available, we intend to facilitate these and other bespoke modifications. Our intention is for the research community to adapt and build upon the OpenIrisDPI platform, tailoring it to the unique demands of their experimental paradigms and advancing the capabilities of high-precision eye tracking.

## Acknowledgments

This work was funded by NIH OD010425, NIH EY032900 to Gregory D. Horwitz, NIH EY032179 to Jacob L. Yates, and an NSF GRFP fellowship to Ryan A. Ressmeyer.

## Declaration of generative AI an AI-assisted technologies in the writing process

During the preparation of this work, the authors used Claude.ai for revisions. After using this tools, the authors reviewed and edited the content as need and take full responsibility for the content of the published article.

## Appendix A. Defining accuracy, precision, and trueness for eye tracking

Throughout this paper, we adopt the definitions for accuracy, precision, and trueness outlined by the International Organization for Standardization (*Accuracy (trueness and precision) of measurement methods and results — Part 1: General principles and definitions* 2023). Under this definition, the accuracy of a measurement refers to its total deviation from a true value of interest. Accuracy is then decomposed into two parts: precision and trueness. Precision refers to the tendency for repeated measurements to be similar to one another, whereas trueness refers to the closeness of the expectation of any given measurement to the true value of interest. With these definitions in hand, we define how precision and trueness relate to eye tracking systems as follows.

The true value of interest for head-centric eye tracking is typically either the orientation of the globe or the direction of gaze. Though related, these two values are not identical. A full description of the globe’s orientation requires knowing three rotational components (pitch, yaw, and roll), whereas just two components are required to uniquely define the ray projecting from the preferred retinal locus (elevation and azimuth). In this study, we ignore torsional eye motion and only estimate the direction of gaze. ’Trueness’ in this context describes the accuracy of calibration, or, equivalently, the accuracy of the function mapping from eye tracker readings to gaze vectors. ’Precision’ then refers to the ability of an eye tracker to produce similar readings given a fixed true gaze vector. The analyses presented in this study are primarily concerned with quantifying the precision of the OpenIrisDPI system under realistic experimental conditions (though the analysis of LGN neural recordings ultimately depends on the system’s overall accuracy).

However, quantifying precision in vivo is fundamentally complicated by the fact that the eyes of living subjects are never still, even during fixation. Because the eye cannot be held perfectly steady, many metrics have been developed that aim to quantify imprecision during experiments (Blignaut and Beelders 2012). However, these methods often conflate fixational motion with imprecision (as in the bivariate contour ellipse), or may ignore low-frequency sources of imprecision (as is true for the sample-to-sample RMS). In this study, we address these issues by explicitly stating our assumption that measurement noise is additive and white to isolate this noise from other sources.

## Appendix B. The OpenIrisDPI feature tracking algorithm

Pseudocode for OpenIrisDPI’s tracking algorithm is given below in Algorithm 1. The algorithm first detects the pupil boundary by computing the convex hull of all points on the edge of the region with pixels below user-defined threshold. An ellipse is then fit to the identified points to estimate the pupil’s center, width, height, and major axis angle. Next, the approximate location of the CR, which is usually the brightest point in the image, is computed as the centroid of pixels above a threshold set close to the sensor’s saturation level. This coarse estimate is then refined by recomputing the thresholded-rectified centroid of a local region around the estimated center. We use the threshold-rectified centroid because we found it to be faster and more accurate in simulations than the radially symmetric center algorithm for reasonable thresholds (Fig. B.1). Note that P2 can shift the calculated center of mass of the CR, but these shifts are a deterministic function of viewing angle and can be compensated for by calibration (see faint reflection above the CR in figures 1 & 2). Finally, the approximate position of P4 is found by identifying the highest amplitude pixel within the pupil, excluding a region around the CR. The sub-pixel location of P4 is then found by computing the threshold-rectified centroid in a region surrounding the approximate center.

### Algorithm 1

OpenIrisDPI Algorithm.

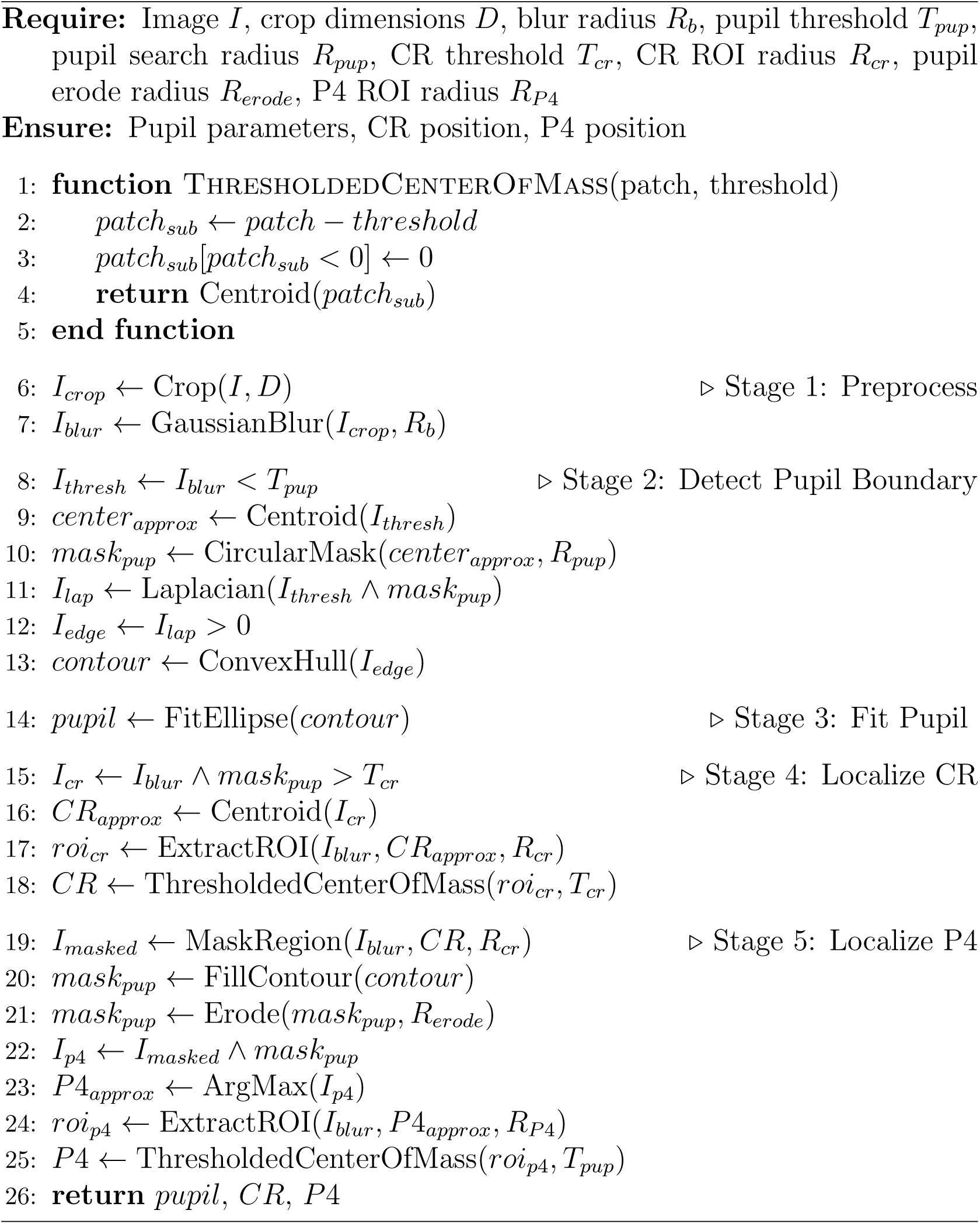

**Figure B.1:**
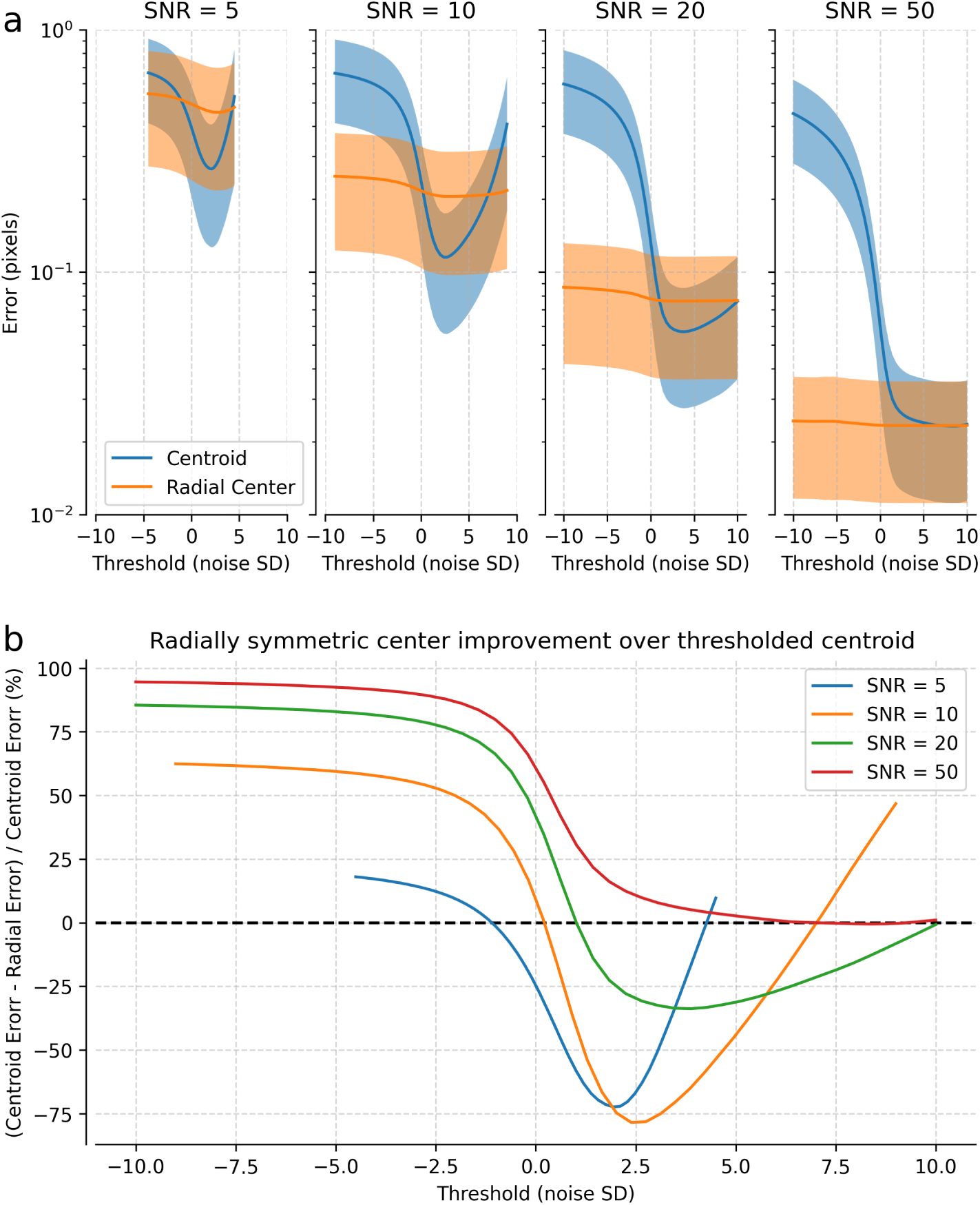
Simulations of localization error using thresholded centroid and radially symmetric center algorithms. a) Localization error across different signal-to-noise ratios (SNR = 5, 10, 20, and 50). Images were simulated as 20 x 20 pixel Gaussian blobs with additive noise 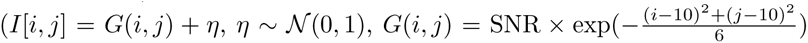). Localization error was defined as the Euclidean distance between the estimated and true center of *G*(*i, j*). Lines show mean ± std of localization error across 10,000 simulated images as a function of threshold value for the thresholded centroid (blue) and radially symmetric center (orange). b) Relative performance difference between methods, normalized by thresholded centroid error. At optimal threshold values, the radially symmetric center is as or more error prone than the thresholded centroid.

## Notes

### Competing Interest Statement

The authors have declared no competing interest.

### Summary of Updates

Introduction revised; Added quantitative safety analysis and irradiance measurements; Added characterization of processing time and latency; Figures 3, 4, and 5 updated to include new panels and annotations; Citation style updated to author-date format.

